# Decidual natural killer cells promote extravillous trophoblast developmental pathways: evidence from trophoblast organoid co-cultures

**DOI:** 10.1101/2024.09.08.611877

**Authors:** Morgan L. Zych, Natalie Lo, Kate A. Patton, Kewei Wang, Brian Cox

**Affiliations:** Department of Physiology, University of Toronto, Ontario, M5S 1A8, Canada

**Author notes:** Authors contributed equally (Natalie Lo, Kate A. Patton, Kewei Wang).

## Abstract

The placenta an essential extraembryonic organ that supports the fetus throughout gestation. The interactions between the placenta and the maternal immune system during the first trimester have not been wholly characterized despite their close physical association and hemi-allogeneic relationship. The most abundant type of immune cell in the uterus in the first trimester is the decidual natural killer cell (dNK). Despite their name, dNKs play supportive roles during pregnancy by remodelling uterine spiral arteries. We present evidence suggesting that the matrix metalloproteinases (MMPs) that dNKs secrete to promote this remodelling also drive placental development. This study used a novel co-culture system of dNKs and trophoblast organoids, which are mini-organs representing two to three different cell types of the human placenta. We found that co-cultures for one week led to significant (p=0.020) increases in the organoid area. We also observed significant decreases in trophoblast stemness markers and upregulation of gene sets associated with extravillous trophoblast (EVT) development through bulk RNA sequencing and immunohistochemical examinations. These changes were accompanied by significant (p<0.001) increases in collagen subunit gene expression in the organoids, with simultaneous significant decreases (p<0.001) in the proportion of organoid area occupied by collagen as determined through Massons Trichrome. Cultures containing dNKs also contained significantly higher MMP1, 3, 9, and 10 levels in their culture media, each of which can break down collagen. These findings demonstrate that dNKs promote changes concordant with trophoblast differentiation towards EVTs and villous branching morphogenesis.

## INTRODUCTION

The placenta is an indispensable organ in mammalian pregnancy. Defects in human placentation can lead to miscarriage or to serious disorders of pregnancy including preeclampsia and fetal growth restriction (Brosens et al., 2011), yet many mechanisms of healthy placentation are not well-understood (Mercuri and Cox, 2022). Trophoblast are the major functional cells of the placenta with respect to nutrient and waste exchange, barrier function, and communication with maternal systems (Knöfler et al., 2019). In human first trimester placental villi, cytotrophoblast (CTB) progenitor cells can differentiate either into multinucleated syncytiotrophoblast (STB) which coat the villi, or extravillous trophoblast (EVT) which migrate away from the placental villi to invade the maternal decidua (Knöfler et al., 2019). As CTB differentiate into EVT, an intermediate cell type called column cytotrophoblast (CCTB) can be identified which is located along the anchoring villi that contact the decidua (Knöfler et al., 2019). The cues that promote CTB transition to CCTB, and CCTB transition to EVT, remain incompletely examined.

The human placenta differs markedly in architecture, immune interactions, and invasion depth from other mammalian placentas, complicating the use of model organisms for its study (Schmidt et al., 2015). Human placental cell lines and explant cultures have considerable limitations as well (Apps et al., 2009; Lee et al., 2016), which researchers have attempted to circumvent with recent culture constructs including human trophoblast stem cells (Okae et al., 2018) and trophoblast organoids. The first human trophoblast organoid protocols were published in 2018 (Haider et al., 2018; Turco et al., 2018) and described methods of deriving these organoids from first trimester primary placental samples. The organoids in each of these protocols are typically described as “inside-out” or “CTB-out” compared to placental villi *in vivo* because they have outer layers of CTB that encase an STB core. Trophoblast organoids have since been derived from blastocysts and from naïve human pluripotent stem cells (Karvas et al., 2022), and have been applied for such purposes as modelling viral entry and screening xenobiotics (Hori et al., 2024; Karvas et al., 2022). Much effort has been made to create trophoblast organoids that are “STB-out” to aid their applications, though this usually employs methods that limit their renewability (Hori et al., 2024). For the present study our group took a different approach and sought to exploit the CTB shell of typical trophoblast organoids to model interactions at the sites where anchoring villi contact the maternal decidua.

The interactions of the hemi-allogeneic fetal trophoblast and maternal immune cells may appear to present an immunological paradox. EVT present fetal antigens to maternal cells through their specialized human leukocyte antigen (HLA) expression, yet maternal cells know they should not respond to these cells as if they were foreign (Tilburgs et al., 2015). Despite this one type of inaction, the immune cells involved in maternal-fetal interactions are also not quiescent during pregnancy. Decidual natural killer cells (dNKs), which are the most abundant type of immune cell resident in the decidua in the first trimester (Monin et al., 2020), assist with spiral artery remodelling through their matrix metalloproteinase secretion (Hazan et al., 2010; Robson et al., 2012; Smith et al., 2009). dNKs have also been found to support the migratory functions of EVT *in vitro* through their secretions (Hanna et al., 2006). dNKs’ killer inhibitory receptors interact with the HLAs expressed by EVT (Vento-Tormo et al., 2018), and enhancing or impairing these interactions can either protect against or promote preeclampsia, respectively (Kennedy et al., 2016; Xiong et al., 2013). Notably, dNKs have been found to secrete growth factors that supply the fetus by crossing the placenta to reach it (Fu et al., 2017a). This led our group to ask whether dNKs also play a role in the development of villous trophoblast. We explored this question using a co-culture model containing trophoblast organoids and allogeneic dNKs. From doing so, we present the first direct evidence of dNKs promoting the growth and development of villous trophoblast towards the EVT fate.

## RESULTS

### Trophoblast organoids and dNKs show appropriate identity hallmarks

Individual components of our co-culture system were first assessed for their expression of proteins and genes that would confirm their identities. Trophoblast organoids were examined with immunofluorescence and found to co-express cytokeratin 7 (KRT7) and GATA3 (Fig. 1A), each of which have been described as key features for identifying villous trophoblast *in vitro* (Lee et al., 2016). Each of these markers localized appropriately with KRT7 staining appearing in the membrane/cytoplasm and GATA3 appearing in the nuclei of trophoblast (Fig. 1A). Trophoblast organoids were also confirmed to express KRT7 by RT-qPCR and found to express trophoblast-identifying markers TFAP2C, ITGA6 (CTB-associated), and ERVW1 (STB-associated) at appropriate levels compared to first trimester placental samples (Fig. 1B). Furthermore, trophoblast organoids and primary placental villi did not show any expression of the decidual stroma markers PRL and IGFBP1, demonstrating that decidua did not contaminate either sample type (Fig. 1B). This RT-qPCR was performed on organoids derived from 6 donors, primary placenta derived from 4 donors, and one decidua donor. All organoids used in subsequent experiments were obtained from our group’s established organoid lines which we cultured, cryopreserved, and thawed extensively, during which time they underwent at least six passages. As such, we concluded that screening these organoids for placental fibroblast markers was not warranted, as fibroblasts were not expected to persist after undergoing these cell divisions.

**Fig. 1.**
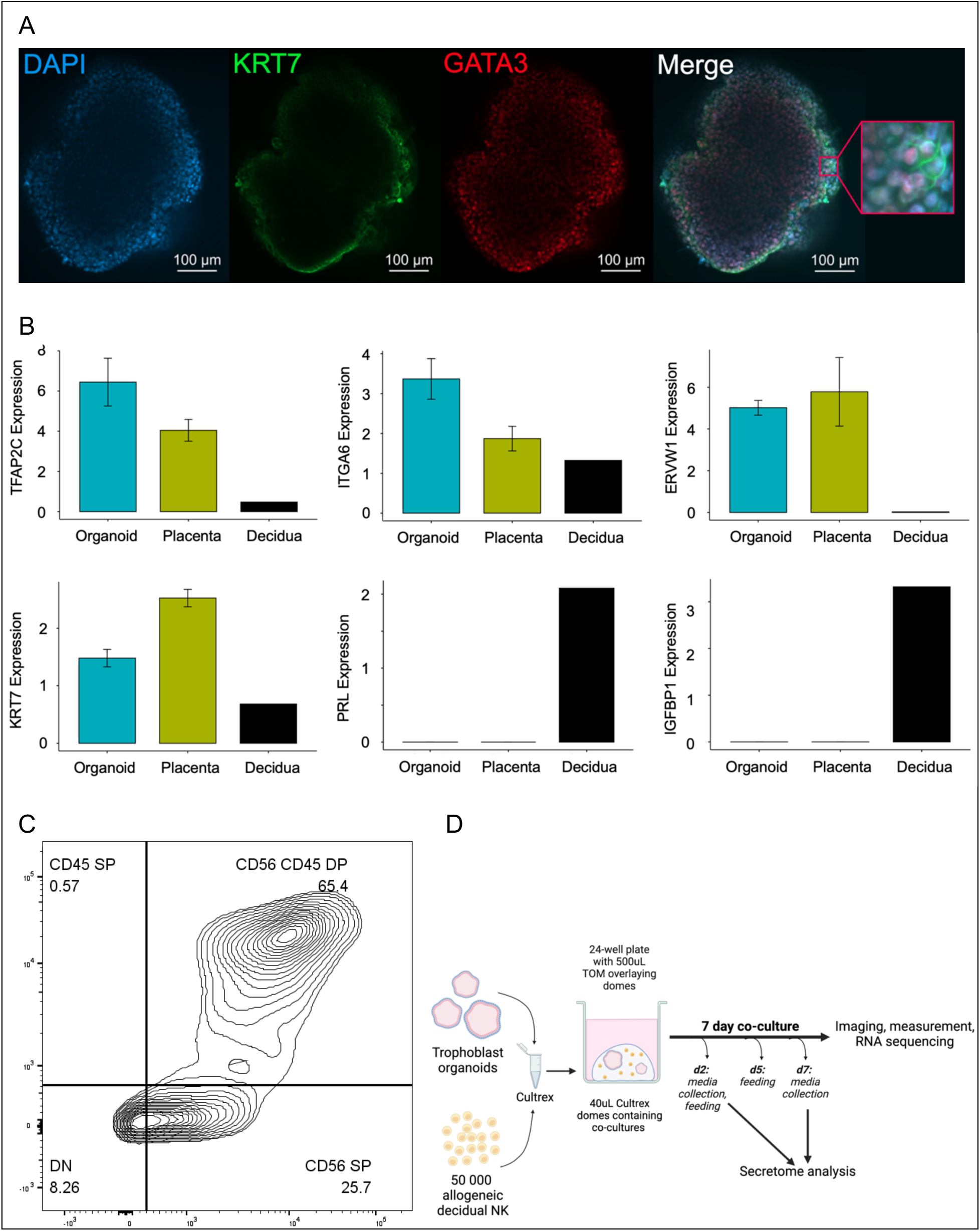
Characteristics of individual culture components. **A.** Trophoblast organoids co-express properly-localized trophoblast markers KRT7 (cell membrane) and GATA3 (nuclear) as shown by immunofluorescent staining. Blue: DAPI counterstaining, green: KRT7 visualized with Alexafluor488, red: GATA3 visualized with Alexafluor647, rightmost image: merge. Inset = additional 5X magnified. **B.** Trophoblast organoids and primary placentas express trophoblast markers TFAP2C, ITGA6, ERVW1, and KRT7 as shown by RT-qPCR. Trophoblast organoids and primary placentas do not express decidua stromal markers PRL and IGFBP1. **C.** Populations of decidual cells present after performing dNK enrichment. Samples from ten donors were stained for CD56 and CD45 expression and analyzed by flow cytometry (SP = single positive, DP = double positive, DN = double negative). Figure shows concatenated results of all ten samples. 91.1% of cells were CD56 positive, with 65.4% of cells double positive for CD56 and CD45. **D.** Culture setup for all subsequent experiments described, created with BioRender.com

While cultures were otherwise performed following the protocols and media formulations described by Turco et al., 2018, we made one key modification and performed them at 2% oxygen. We felt this was essential because the oxygen level measured at the maternal-fetal interface before spiral artery remodelling is 2% (Jauniaux et al., 2000; Rodesch et al., 1992), and because hypoxia promotes tolerogenic signalling and dNK-associated phenotypes in both human and murine NK cells (Cerdeira et al., 2013; Kenchegowda et al., 2017; Parodi et al., 2018). Flow cytometry was performed on decidual cells from 10 donors following enrichment for dNKs. This analysis was concatenated in the plot shown in Fig. 1C and shows that 91.1% of live cells analysed were positive for the canonical NK marker CD56. Variable levels of CD45 were detected in these cells (Fig. 1C), as is expected in tissue-resident CD56^Bright^ populations (Krzywinska et al., 2016). Details of individual donor measurements and gating strategies can be found in Fig. S1A-B. From these analyses, we considered this cell population to be sufficiently enriched for dNKs, and elected to move forward with their co-culture with trophoblast organoids. A summary figure of the co-culture setup and key time points for all analyses which followed is provided in Fig. 1D. Briefly, trophoblast organoids were co-cultured with allogeneic dNK cells by seeding both together into the same extracellular matrix (ECM) gel domes (Fig. 1D). Trophoblast organoid media was then overlaid on top of polymerized gel domes (Fig. 1D). Co-cultures and monoculture controls for both organoids and dNKs then proceeded for 7 days with media exchange occurring on days 2 and 5 (Fig. 1D). Key to the design of all subsequent experiments was the inclusion of organoids and dNKs that were from a minimum of two allogeneic donors each, cultured together in minimum four combinations. Details of sample sizes, donor representation, and statistical tests used in each subsequent analysis can be found in Supplementary Table 1.

### Morphological changes in dNK-co-cultured trophoblast organoids

To understand changes that occurred to the gross structure of trophoblast organoids as a result of their co-culture with dNKs, monocultured and co-cultured trophoblast organoids were examined through whole mount confocal microscopy. Analysis of 50 monocultured organoids and 82 co-cultured organoids representing nine different donor combinations (four trophoblast and five dNK) showed significant (p<0.05) increases in 2D area of co-cultured organoids compared to their monocultured counterparts after seven days (Fig. 2A). Not all donor tissue combinations responded equally, although a majority of donor combinations show increased organoid area upon co-culture (Fig. S2A). We realized that a possible alternative explanation of our observations could be more frequent new organoid development in monoculture, such that proliferating trophoblast contributed to new small aggregates rather than expanding the area of existing structures. We evaluated this possibility by comparing the total numbers of organoids between mono- and co-culture and observed no significant differences (Fig. S2B), demonstrating that co-culture did not alter the rate of organoid formation.

**Fig. 2.**
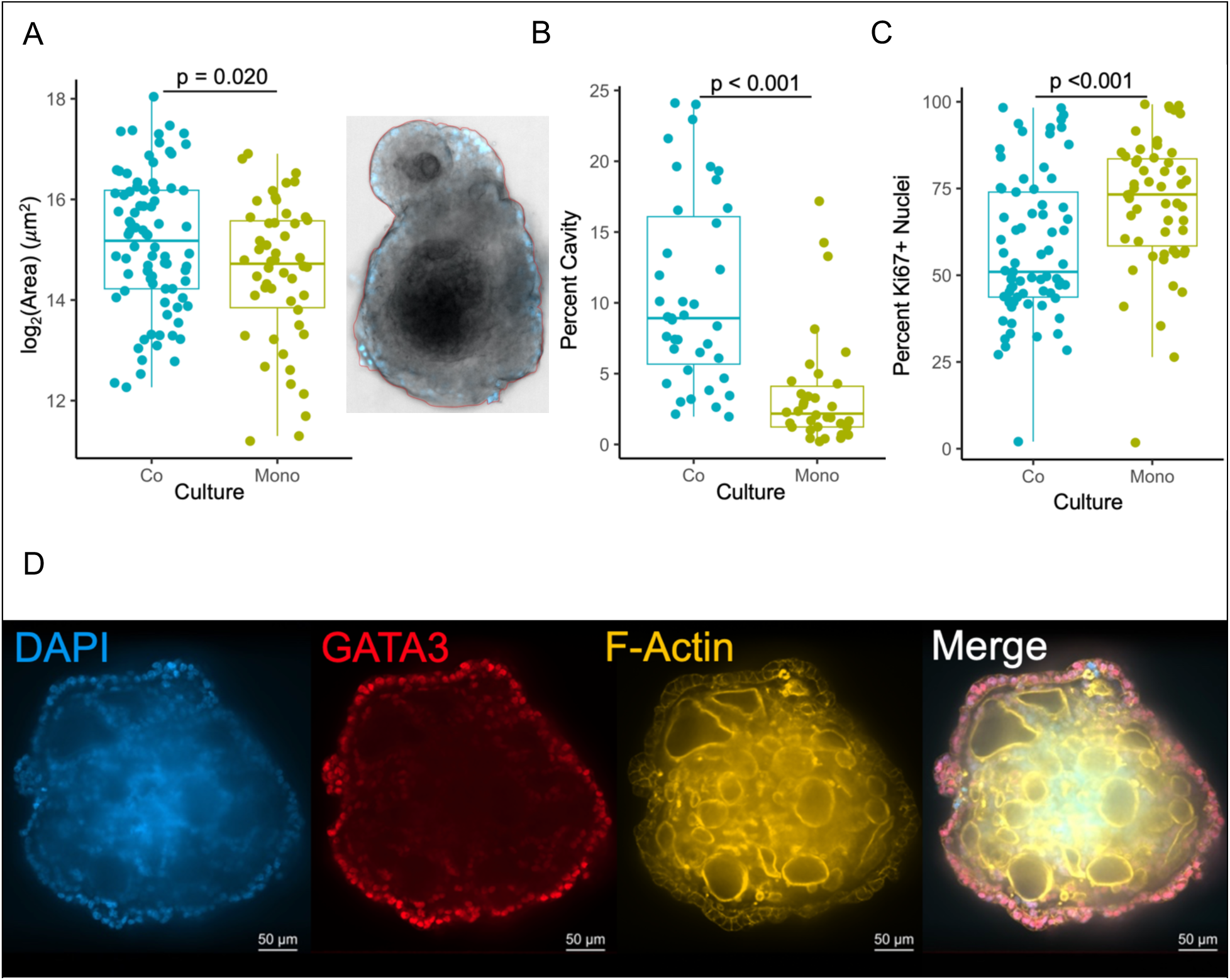
Trophoblast organoids increase in 2D area and in proportion of 2D area occupied by cavities following co-culture with allogeneic dNKs. **A.** Significant increase (p=0.020) in 2D area of trophoblast organoids is observed after co-culture with allogeneic dNKs for 7 days. An example of whole mount bright field image of organoid for prepared 2D area measurement is shown to the right of the boxplot. **B.** Significantly larger proportions of trophoblast organoid area are occupied by cavities after co-culture with dNKs for 7 days (p<0.001). **C.** Significantly reduced proportion of nuclei stained positive for Ki67 (proliferating cells) (p<0.001) in organoids co-cultured with dNKs versus monocultured organoids. **D.** Trophoblast organoids contain cavities within their syncytialized regions bordered by F-actin as shown by immunofluorescent examination of a mono-cultured organoid. Blue: DAPI counterstaining, red: GATA3 visualized with Alexafluor647, orange: F-actin visualized with ActinRed 555 ReadyProbes, rightmost image: merge.

We next hypothesized that the observed size differences between monocultured and co-cultured organoids could be a result of trophoblast proliferation, structural rearrangement of the organoids, or of a combination of both factors. Trophoblast organoids commonly display cavities (Turco et al., 2018), which can manifest as non-cellular regions within sectioned organoids, and as F-actin-bordered empty spaces within whole mount immunofluorescent stained 3D organoids (Fig. 2D). Trophoblast organoids were found to have a significantly larger percentage (p<0.001) of their 2D area occupied by cavities when they had been co-cultured with dNKs for 7 days (Fig. 2B). This comparison was made by measuring the proportion of organoid area occupied by non-cellular regions within 2D sections, ensuring that measurements were made at the middle of each organoid by examining serial sections. This finding suggests that the increase in cavitation drives the overall observed increase in organoid size, at least in part. A reduction in the amount of Ki67+ nuclei was also observed in co-cultured organoids through immunohistochemical staining (Fig. 2C), further demonstrating that morphological changes rather than proliferation drive observed organoid size increases. To better characterize the mechanisms behind these observed structural changes, we went on to analyze organoid transcription, marker protein expression, and secreted factors in culture media.

### Bulk transcriptomic analysis of trophoblast organoids, dNKs, and their co-cultures

To examine which transcriptional changes may underlie the observed increase in trophoblast organoid size upon co-culture with dNKs, we performed bulk RNA sequencing on monocultured and co-cultured organoids, and on monocultured dNKs. A principal components plot was prepared showing gene expression of trophoblast organoids from two donors, dNKs from two donors, and all four combinations of these constituents in co-culture (Fig. 3A). Batch correction proceeded to adjust for multiple sequencing runs and differences in sample preparation. dNK monocultures separate from all organoid-containing cultures when examined by their principal components, while individual dNK samples show considerable distance from each other as well (Fig. 3A). Samples from organoid monocultures and co-cultures are somewhat interspersed, but are still broadly distinguishable (Fig. 3A). Differences in sample preparation occurred due to attempts to separate organoids and dNKs following their co-culture by spinning them on a Ficoll gradient. Cultures that were subject to this treatment clustered together with cultures that did not undergo this treatment in their principal components (Fig. 3A), suggesting that Ficoll treatment had minimal impact on dNK removal. To examine differences in expression that were specific to trophoblast organoids, we used statistical methods to remove dNK-associated expression from all co-cultured organoid samples (Fig. S3).

**Fig. 3.**
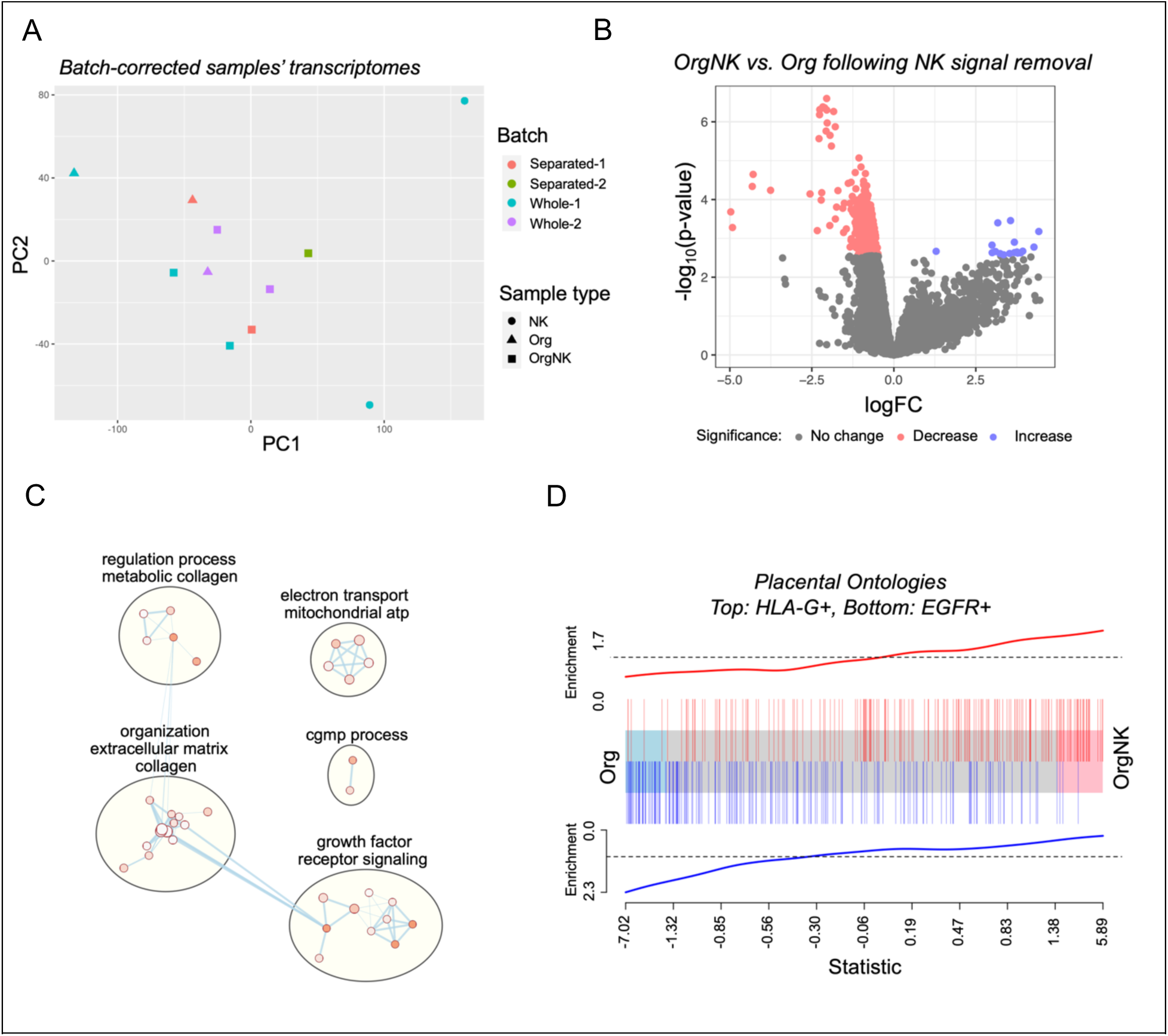
dNKs prompt transcriptional shifts in trophoblast organoids towards extracellular matrix remodelling and EVT development. **A.** Principal components plot derived from bulk RNA sequencing of trophoblast organoids, dNKs, and their co-culture. **B.** Volcano plots of differentially expressed genes between organoids in co-culture versus monoculture both with and without correction for dNK expression. **C.** Cytoscape Plot of the most significantly differing gene ontologies found in trophoblast organoids in co-culture versus monoculture. D. Barcode plot prepared using Placental Gene Sets showing enrichment of HLA-G+ EVT-related gene sets and reduced EGFR+ CTB-associated gene sets are expressed in organoids co-cultured with dNKs compared with monocultured organoids.

Once the statistical filtering step removed confounding dNK signals (Fig. S3), monocultured and co-cultured organoids showed multiple significantly differentially expressed genes (Fig. 3B). This was explored further by assessing the Gene Ontologies of differentially expressed genes and displaying them in Cytoscape. The Gene Ontologies that differed between monocultured and co-cultured trophoblast organoids were mainly related to collagen (Fig. 3C) with respect to both its production and its organization. ‘Growth factor receptor signalling’ was also a broad category of ontology that arose, suggesting that organoids may be responding to secreted signals from dNKs to prompt them to grow and develop. Additionally, some energy usage-related ontologies were found to be important, with these being characterized as ‘electron transport mitochondrial ATP” by Cytoscape. Perhaps these ontologies arose in response to increased energy needs and shifting metabolism as organoids rearrange their gross structures and as cells within them differentiate. Finally, two ontologies related to ‘cGMP process’ were identified, but the implications of these are unclear.

Our group has previously identified a limited representation of placenta-related gene sets within the general Gene Ontologies analysis list (Naismith and Cox, 2021). To account for this under-representation, a separate analysis was performed with gene sets prepared specifically from placenta and placenta-associated datasets termed Placental Gene Sets (Fig. 3D) (Naismith and Cox, 2021). This analysis clearly demonstrated an enrichment of EVT differentiation-associated gene sets at the expense of EGFR-associated (CTB-associated) gene sets in trophoblast organoids co-cultured with dNKs (Fig. 3D). These findings suggest that the observed increases in trophoblast organoid size and the structural rearrangements that they undergo when co-cultured with dNKs (Fig. 2A-B) are accompanied by increased differentiation of CTB towards the EVT fate, and by increased extracellular matrix remodelling that is expected upon cells undergoing epithelial-to-mesenchyme transition (EMT). This conclusion was supported by plotting the observed Placental Gene Sets in Cytoscape (Fig. S4A). We next quantified the expression of differentiation-associated proteins and collagen staining in trophoblast organoids to attempt to validate our conclusions from our transcriptomic analysis.

### Visualization of differentiation-related proteins and collagen deposition within trophoblast organoids

Our transcriptomic data strongly suggested that co-culture of trophoblast organoids with dNKs promotes trophoblast differentiation along the EVT pathway. In addition to the observed size increases upon co-culture, the organoids retained their globular structures. These observations together suggest that co-culture with dNKs does not lead to mature EVT development. Development of mature EVTs in this culture context leads to their migration away from the source organoid, as is observed when organoids are cultured in EVT-promoting media (Turco et al., 2018). Furthermore, HLA-G, the canonical marker for EVTs, did not show differential expression in our transcriptomic analysis (Fig. S4B). Consequently, we looked to validate that differentiation towards the EVT fate was occurring using a different EVT-associated marker, CD44, by evaluating its protein expression through immunohistochemistry. A significantly greater proportion of organoids expressed CD44 protein when co-cultured with dNKs (p<0.001) (Fig. 4A).

**Fig. 4.**
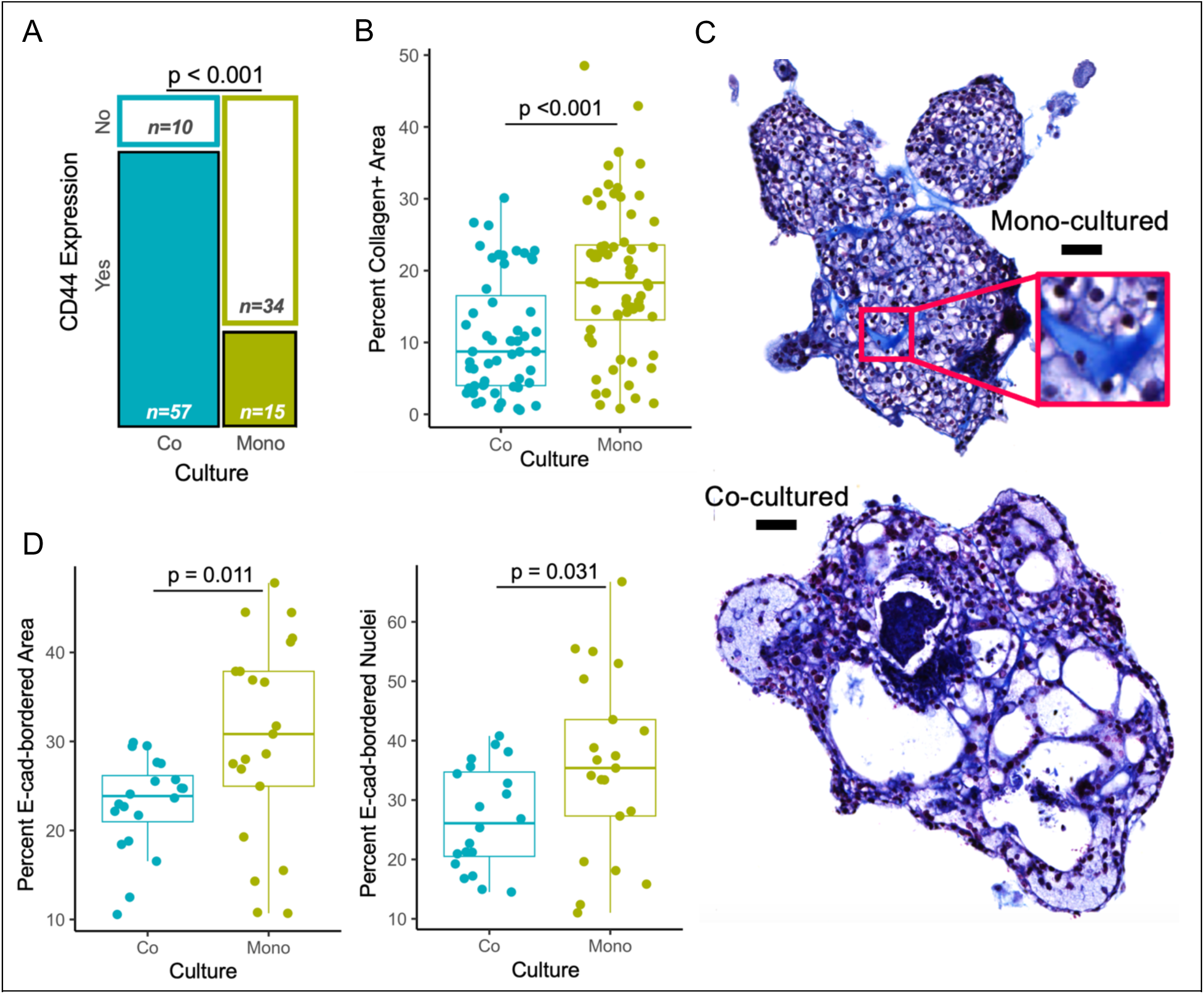
Protein expression and localization congruent with villous maturation is observed in trophoblast organoids upon co-culture with allogeneic dNKs. A. Significantly increased proportion of organoids stain positive for the EVT marker CD44 upon co-culture with dNKs (p<0.001). B. Significant differences in Masson’s trichrome staining between monocultured trophoblast organoids versus those co-cultured with dNKs, as assessed by area measurements. C. Examples of Masson’s trichrome staining to visualize collagen in cultures (bright blue, scale bar = 50μm, inset zoom = 2.5x). D. Significantly reduced proportion of organoid area bordered by cell membrane stained positive for E-cadherin (CTB) (p=0.011) and in proportion of nuclei bounded by cell membrane stained for E-cadherin (p=0.031) in organoids co-cultured with dNKs versus monocultured organoids.

Multiple collagen-related gene ontologies appeared in our analysis of the differentially expressed genes of co-cultured organoids. To better understand the impacts of these changes on the spatial organization of collagen in trophoblast organoids, we performed Masson’s trichrome staining (Fig. 4B-C), which in these samples results in purple cytoplasm, dark pink nuclei, and collagen of all types stained bright blue. Collagen staining was visualized using the blue chromaticity function on QuPath for quantification with both point counting (overlaying a grid onto images and categorizing each resulting intersection point of the gridlines as either collagen-positive or -negative) and with AI-guided pixel classification. Despite observing increased expression of genes for multiple collagen fibre subunits in co-cultured organoids, the proportion of organoid area occupied by collagen staining was found to significantly decrease in co-cultured organoids (p<0.001) (Fig. 4D). This change was not accompanied by any differences in distribution of collagen, as there was no difference in the proportion of collagen-positive points found in monocultured and co-cultured organoids (Fig. S5A).

Next, we looked to validate that trophoblast was differentiating in co-cultured organoids by performing IHC for E-cadherin, hypothesizing that its levels would be reduced upon co-culture. Both total area and the proportion of nuclei occupied by E-cadherin-bordered staining in trophoblast organoid cultures were significantly reduced upon their co-culture with dNKs (p<0.05, p<0.05) (Fig. 4A). In trophoblast, E-cadherin serves as a CTB marker, the loss of which reflects differentiation as well as a shift in the cell-cell interactions that E-cadherin is known to mediate. With this additional evidence that trophoblast within co-cultured organoids is differentiating, we also examined the possibility that the STB was altered upon co-culture. Proportions of organoid area bordered by the STB marker SDC1, and proportions of nuclei contained within these stained regions both remained unchanged upon co-culture with dNKs (Fig. S5B). To gain insight into possible mechanisms through which dNKs promote EVT-directed development, and to resolve apparent discrepancies in our findings regarding collagen deposition, we next explored changes to the secretome of trophoblast organoids, dNKs, and their co-cultures.

### Secreted factors in media of trophoblast organoids, dNKs, and their co-cultures show dNKs drive villous remodelling and development

Media from trophoblast organoid monocultures, dNK monocultures, and their combinations were analyzed using two different panel assays to determine how secreted factors contribute to trophoblast-dNK crosstalk. The first panel assayed 48 immunomodulatory factors (Fig. 5A), and the second panel assayed 30 factors involved in angiogenesis, growth, and extracellular matrix remodelling. Media samples were collected on day 2 and day 7 of culture (Fig. 1D). The principal components plot derived from the first panel (Fig. 5A) shows that variance of these immunomodulatory factors in culture supernatants is driven by the presence of dNKs. Conversely, organoid monocultures, dNK monocultures, and their co-cultures can be readily distinguished from each other in the principal components plot prepared from the second panel (Fig. 5B).

**Fig. 5.**
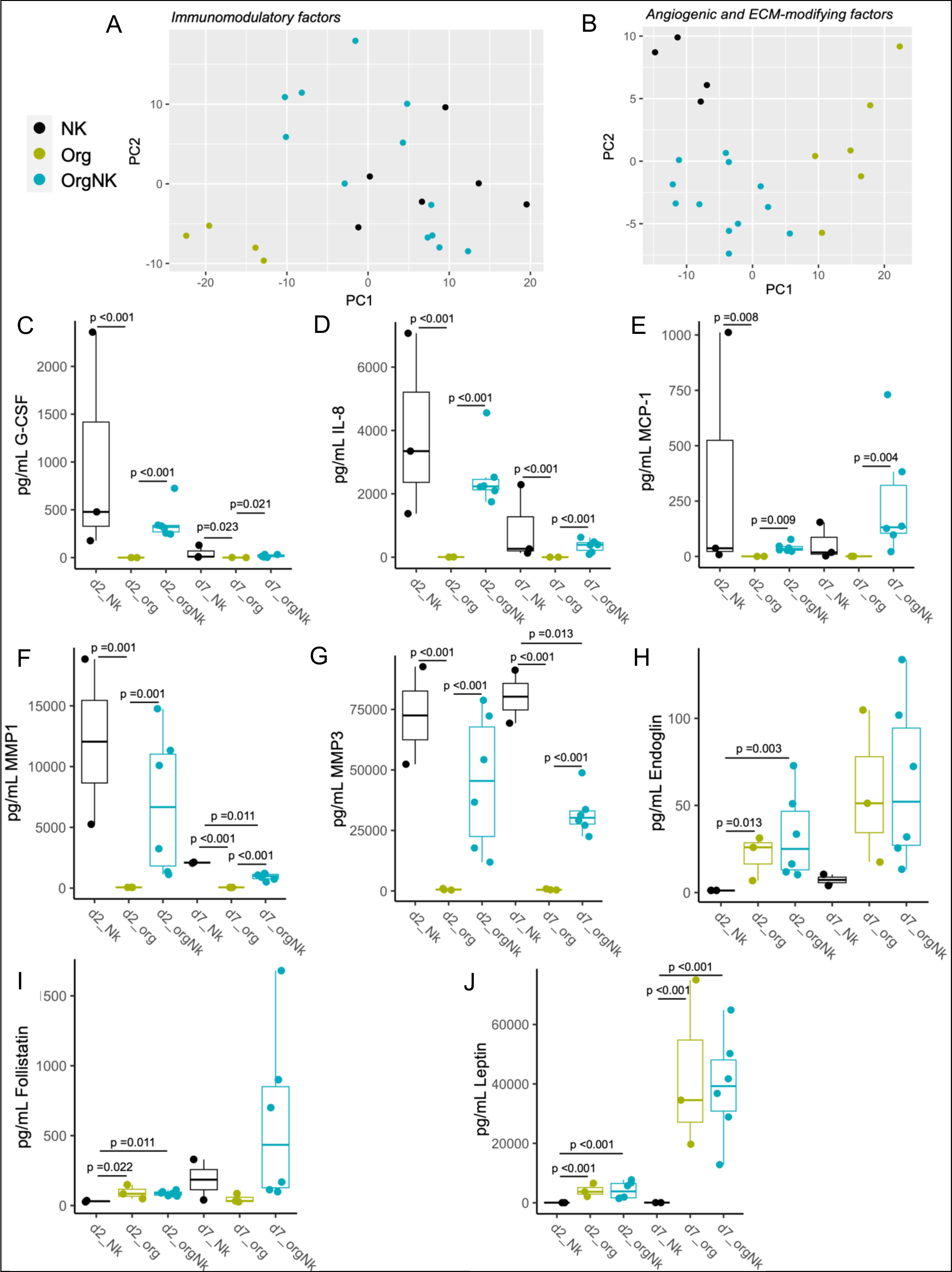
Measurement of immunomodulatory, pro-angiogenic, and ECM-modifying factors in culture media. **A-B.** Legend for the entire figure and principal components plots for media of trophoblast organoid and dNK monocultures, and their co-cultures that was assessed using a panel of 48 immunomodulatory factors (**A**) and a panel of 30 factors related to angiogenesis, extracellular matrix modification, and growth that a separate group of samples was subjected to (**B**). **C-D**. G-CSF (**C**) and IL-8 (**D**) levels are increased in media containing dNKs, with higher levels seen on day 2. **E.** On day 7 of culture, highest levels of MCP-1 are seen in co-culture. **F-G.** MMP1 (**F**) and MMP3 (**G**) levels are increased in media containing dNKs. **H.** Endoglin levels are increased in media containing trophoblast organoids, with highest levels seen on day 7. **I.** Follistatin levels are increased in media containing trophoblast organoids on day 2. While not significant, highest levels are seen in co-culture on day 7. **J.** Leptin levels are increased in media containing trophoblast organoids, with highest levels seen on day 7.

The panel of immunomodulatory factors demonstrated that G-CSF (Fig. 5C), IL-8 (Fig. 5D), and IL-6 (Fig. S6A) were present in dNK-containing cultures. Each of these factors has demonstrated importance at the maternal-fetal interface (Ding et al., 2021; Jovanović et al., 2010; Pitman et al., 2013) and their presence in this culture construct suggests that physiological responses to these signals by trophoblast could be occurring. However, each of these factors was found in smaller quantities on day 7 of culture than on day 2 (Fig. 5C-D, Fig. S6A), suggesting that any signals dNKs provide to trophoblast organoids through these factors may be intermittent, or temporary. The impacts of IL-8 on trophoblast organoid growth were further investigated due to its established role in EVT migration (De Oliveira et al., 2010; Jovanović et al., 2010). Exogenous IL-8, added in the absence of dNKs, was not found to impact 2D organoid area (Fig. S6E). Intriguing trends were observed in amounts of MCP-1 found in culture media, with increased levels in all dNK-containing cultures, and with the highest levels on day 7 found in co-cultures (Fig. 5E). This finding suggests that while trophoblast organoids and dNK acclimate to each other’s presence over time, MCP-1 production is promoted.

The second panel of secreted factors revealed that dNK-containing cultures have significantly higher levels of MMP1 (Fig. 5F), MMP3 (Fig. 5G), MMP9 (Fig. S6B), and MMP10 (Fig. S6C) than trophoblast organoid monocultures. The presence of these factors provides clear insight into the observed reduction of collagen-stained area in co-cultured trophoblast organoids (Fig. 4D), as each of these MMPs can digest collagen. The MMP2 gene, a marker of CTB transition to EVT (DaSilva-Arnold et al., 2015), was found to be upregulated in co-cultured trophoblast organoids, however no significant differences were observed in levels of MMP2 present in culture media (Fig. S6D). These findings together with the increased expression of collagen subunit genes and their associated pathway members in co-cultured trophoblast organoids suggests that processes of dynamic remodelling, congruent with villous development, are occurring in these organoids.

Secreted factors that likely have trophoblast organoids as their sources were also observed in this panel and these may also contribute to the cross-talk between culture constituents. Endoglin levels are increased in culture media containing trophoblast organoids, with highest levels observed on day 7 (Fig. 5H). On day 2, follistatin levels are increased in media containing trophoblast organoids, but on day 7 its levels are reduced in trophoblast organoid monocultures, while it is still increased in co-cultures (Fig. 5I). Leptin levels were found to be increased in all trophoblast organoid-containing cultures as well, with its highest levels observed in co-cultures on day 7 (Fig. 5J). While levels of organoid-derived factors vary considerably, likely as a result of differences in the populations of secreting trophoblast between each culture preparation, these factors could still represent influences on, or results of, dNK-trophoblast communication.

## DISCUSSION

Previous work exploring the cross-talk between dNKs and trophoblast has primarily focused on interactions between dNKs and fully developed EVTs. These cell types have been found to collaborate for the purposes of chemotaxis and colocalization (Hazan et al., 2010), spiral artery remodelling through extracellular matrix degeneration (Choudhury et al., 2019), and immunoregulation of the cells present in the decidua (Vento-Tormo et al., 2018). The development of EVTs from villous trophoblast precursors has been described as “decidua-independent” due to the occurrence of spontaneous EVT growth from villous explant cultures (Pollheimer et al., 2018). In contrast, our group has uncovered evidence of cross-talk which occurs during trophoblast development, wherein dNKs play a role in promoting EVT differentiation from villous trophoblast precursors.

The relationship between disorders of pregnancy with both shallow placentation (Brosens et al., 2011) and improper trophoblast-dNK communication (Zhang and Wei, 2021) is well-described. Recurrent pregnancy loss has been associated with reduced CD49a expression on dNKs leading to reduced tolerance (Li et al., 2019) , as well as reduced proportions of a dNK subtype associated with angiogenesis and interaction with EVT (Guo et al., 2021). While preeclampsia manifests in heterogeneous ways (Leavey et al., 2016), one of the most consistent phenomena associated with this serious disorder is immune dysfunction, including dNK dysfunction (Fu et al., 2023; Leavey et al., 2019). Genetic contributions to preeclampsia are similarly heterogeneous, but one of the most consistently documented risk allele combinations is maternal dNK KIR2DL1 and fetal HLA-C2 (Hiby et al., 2004; Xiong et al., 2013). Similar communication axes between dNKs’ KIR and trophoblast’s EVT have been found to impact birth weight in humans (Hiby et al., 2014), and dNK population deficits are reliably associated with growth restriction in mice (Boulenouar et al., 2016; Fu et al., 2017a). Our co-culture model provides a tunable system where the dNK dysfunction phenotypes associated with disorders of pregnancy can be assessed for their effects on upstream villous trophoblast development in the future. Improved understanding of the biomolecules, dNK-derived or otherwise, that we expect to find at the maternal-fetal interface at the first trimester is a promising avenue for developing biomarkers and therapeutics for disorders of pregnancy. In our trophoblast organoid-dNK co-cultures, we observed various secreted molecules including multiple MMPs, cytokines, chemokines, and hormones that are present at the first trimester maternal-fetal interface (Caniggia et al., 1997; Choudhury et al., 2019; Ding et al., 2021; Jovanović et al., 2010; Li et al., 2022; Pitman et al., 2013; Qin et al., 2023; Toro et al., 2014; Vettraino et al., 1996). These observations support the physiological relevance of our co-culture model and also provide clear directions for future work, as the roles of each of these secreted factors in promoting trophoblast function is somewhat established, but their roles in driving different trophoblast developmental fates are less clear.

Our findings align well with those of Li et al, whose group cultured trophoblast organoids with cytokines that they identified as being key to trophoblast-dNK interactions. This group found that application of these cytokines to trophoblast organoids also increased their propensity to differentiate towards the EVT fate (Li et al., 2024) Our panel of immunomodulatory factors analysed in culture media did not capture any of the cytokines (CSF1, CSF2, XCL1, and CCL5) that this group applied to organoids. Future work to determine whether these cytokines are present in trophoblast organoid co-cultures with dNKs and how these cytokines impact trophoblast organoid morphology and their ECM is sure to provide fascinating insights.

Some markers of column CTB and EVT differentiation, most notably HLA-G, did not show significant expression differences in co-cultured organoids. These results are likely a reflection of the heterogeneous states of the trophoblast represented in the organoids, with a continuum of those differentiating towards the EVT fate present. Complete transition to fully-realized EVTs in this context also may not have been possible given the Wnt-activating components of trophoblast organoid media (Haider et al., 2018). The markers which did show significant differences (ASCL2, FOXO1, CD44) are some of those that have been shown to play roles specifically in the invasiveness of EVTs (Chen et al., 2018; Takahashi et al., 2014; Varberg et al., 2021). The ability of dNKs to enhance the invasive properties of developed EVTs has been established in various culture settings (Hanna et al., 2006; Mani et al., 2024). However, there is some suggestion that it is only dNKs from later in gestation than those we examined that would be expected to enhance EVT invasion (Lash et al., 2010). The findings of the present study that describe a role for dNKs in prompting EVT differentiation may help explain the gestational age-dependent differences observed in previous work involving mature EVTs.

Beyond serving as a marker for EVT fate, our finding of CD44 gene expression as well as protein upregulation in co-cultured organoids may provide insight into the mechanisms of dNK-driven organoid remodelling as well. The invasive capability of EVT has been shown to depend on their CD44 expression, the ligand of which is ECM component hyaluronic acid (Takahashi et al., 2014). It is unsurprising that we observed differentiation of trophoblast along the EVT pathway concomitant with alterations to the extracellular matrix given the association of EVT differentiation with epithelial-to-mesenchyme transition (DaSilva-Arnold et al., 2015). The process of epithelial-to-mesenchyme transition varies based on its context but consistently involves loss of epithelial markers (such as E-cadherin) as well as ECM changes, both of which were observed in our trophoblast organoid co-culture (DaSilva-Arnold et al., 2015). Primary EVT have also been observed to express collagen genes (Oefner et al., 2015), a phenomenon which was validated in our gene expression data of co-cultured organoids. Additionally, trophoblast-secreted collagen has been found to promote tolerogenic dNK activities (Fu et al., 2014; Fu et al., 2017b). The collagen subunit gene expression we observed, together with matrix metalloproteinase expression, and the net reduction in collagen surface area in co-cultured organoids suggests that processes of dynamic remodelling are occurring in these contexts.

One key limitation of this study was our performance of extracellular matrix analysis in cultures that were performed within basement membrane extract gel, which had the potential to confound our findings. Future work involving trophoblast organoid and dNK co-culture should be performed using culture methods that do not require ECM gel domes, to better uncover the changes to endogenous ECM that are promoted by dNK-trophoblast interactions. The insights that can be gained from co-culture models of the maternal-fetal interface can also be expanded by incorporating additional cell types, including endothelial cells and decidual stromal cells. The scale of the co-cultures performed also did not permit detailed examination of the changes undergone by the dNKs as a result of their culture with trophoblast. This side of the “conversation” between trophoblast and dNKs therefore remains under-examined. However, the co-culture construct presented in the present study provides a reproducible and reliable model of this communication axis at the anchoring placental villus for future work.

## METHODS

### Placenta and decidua processing

Placenta and decidual samples were obtained through the Research Centre for Women’s and Infants’ Health at the Lunenfeld-Tanenbaum Research Institute, Mount Sinai Hospital, Toronto. Elective first trimester pregnancy termination patients provided informed consent to donate their tissues, and collection proceeded according to Research Ethics Board-approved protocols. Placental tissues dating from 4.0 to 7.5 weeks of gestation were processed for trophoblast organoid derivation by dissection into 1mm x 1mm pieces. Placenta fragments were processed using a series of three 10-minute digestions in trypsin buffer solution, followed by application of 10% FBS (Gibco A3160702) in HBSS to stop the reaction, passage through a 100μm mesh filter, and washes between each digestion. Each digestion step took place on a benchtop shaker set to 250rpm and 37°C. Each 50mL of trypsin buffer solution contained 5mL of 10X Trypsin (Gibco 15090046), 0.5mL Antibiotic-Antimycotic (Gibco 15240062), 100μL DNAse I (prepared at 100U/μL with Roche 10104159001), and 44.4mL digestion buffer (each 50mL of digestion buffer contained 1.25mL HEPES (BioShop HEP003) and 0.21mL Mg_2_SO_4_ (Boston Bioproducts MT-210) in HBSS (Gibco 14175095)). Isolated trophoblast cells were then cryopreserved in aliquots of <2x10^6^ cells/mL freezing medium (10% DMSO in FBS). Decidual tissues dating from 4.0 to 12.0 weeks of gestation were processed for dNK selection by dissection into 1mm x 1mm pieces.

Decidua fragments were processed by digesting them in 10mL Collagenase IV buffer solution with 20μL DNAse I per sample. Collagenase IV buffer solution was comprised of 500mg powdered Collagenase IV (Gibco 17104019), 200mL HBSS, 22.3mL FBS, and 1190μL bovine serum albumin solution (Sigma-Aldrich A9576). These fragments were digested for 75 minutes on a benchtop shaker set to 250rpm and 37°C. Isolated cells were passed through a 100μm mesh filter, washed, applied to a Ficoll Paque Plus (Cytiva 17-1440-03) gradient, and spun according to the manufacturer’s instructions. Cells from the “buffy coat”, which are enriched for viable lymphocytes, were then subjected to magnetic bead-based NK cell negative selection using the EasySep Human NK Enrichment Kit (Stemcell Technologies 19055). Selected cells were cryopreserved in aliquots of <2x10^6^ cells/mL freezing medium (10% DMSO in FBS).

### Trophoblast organoid culture

Trophoblast organoids were cultured according to the protocol described by (Turco et al., 2018), with the following modifications: mouse HGF (Sino Biological 50038-MNAH) rather than human HGF was used, cultures were performed using reduced growth factor Cultrex basement membrane extract (R&D #3433), and cultures were maintained in hypoxia chambers (Billups-Rothenberg MIC-101) perfused with 2% O2, 5% CO2 for 5 minutes at a rate of 25 L/min. Media was pre-conditioned in the perfused hypoxia chamber in a 37°C incubator for one hour before being applied to cultures. Trophoblast organoid media was comprised of: 1X N2 supplement (Gibco 17502048), 1X B27 supplement minus vitamin A (Gibco 12587010), 2mM L-glutamine (R&D B90010), 1.25 mM N-Acetyl-L-Cysteine (Cayman Chemical 20261), 50ng/mL human EGF (Gibco PHG0311L), 100ng/mL human FGF2 (Sigma-Aldrich F0291), 50ng/mL mouse HGF (Sino Biological 50038-MNAH), 80ng/mL human R-spondin (R&D 4645-RS), 500nM A83-01 (Tocris 2939), 2.5uM prostaglandin E2 (Tocris 2296), 1.5uM CHIR99021 (Cell Signaling Technology #54290S), 2uM Y27632 (Cell Signaling Technology #13624), and 1X Antibiotic-Antimycotic (Gibco 15240062) added to Advanced DMEM/F12 (Gibco 12634010). Trophoblast organoid cultures were initiated by seeding each dome of Cultrex gel either with <50 000 of thawed primary trophoblast cells or with thawed cryopreserved organoids from previous cultures. 40μL domes of Cultrex gel containing cultures were prepared in 24-well plates, allowed to polymerize upside-down, and were then overlaid with 500μL of trophoblast organoid media per dome. When co-cultures with dNKs or dNK monocultures were prepared, 50 000 dNKs were added to each 40μL Cultrex dome. This number of dNKs was selected because it was the largest number that could be accommodated before their presence caused the Cultrex gel to depolymerize.

### Whole mount microscopy and Immunofluorescence

Cultures that were subjected to whole mount immunofluoresent imaging were grown on 24-well glass bottom plates (Cellvis P24-1.5H-N). All staining took place within these plates. Organoid area comparisons were made on whole mounted samples counterstained with DAPI (Thermo Scientific 62248). Their imaging was performed on the Zeiss Spinning Disk Confocal AxioObserverZ1 in the University of Toronto’s Microscopy Imaging Laboratory facility to generate optical sections. 2D area at each organoid’s widest point was then measured using Zeiss ZEN software. For immunofluorescent staining, samples were fixed in pre-warmed paraformaldehyde for 15 minutes. Samples were then washed and permeabilized with 0.25% Triton-X-100 (Sigma X100) for 10 minutes. Samples were washed again, then blocked with 1% BSA in PBS for 30 minutes. Details of all antibodies used and their dilutions are described in Supplementary Table 2. Primary and secondary antibodies were each incubated with samples for one hour at room temperature, with washes in between. Samples were then counterstained with DAPI (Thermo Scientific 62248), washed, and imaged in PBS using the Zeiss Spinning Disk Confocal AxioObserverZ1 in the same facility.

### RT-qPCR

Snap frozen tissue and culture samples were placed in TRI Reagent (Sigma-Aldrich T9424) immediately upon removal from the freezer. RNA was isolated according to TRI Reagent manufacturer’s instructions. RNA was quantified using the QuBit High Sensitivity system, and cDNA was prepared using the High Capacity cDNA Reverse Transcription kit from Thermo Fisher Scientific (4368813). For qPCR, TaqMan Gene Expression assays from Applied Biosystems were prepared using TaqMan 2X Universal PCR Master Mix (Thermo Fisher Scientific 4304437). The genes assayed and their assay catalogue numbers are as follows: ITGA6 (Hs01041011), TFAP2C (Hs00231476), KRT7 (Hs00559840), ERVW1 (Hs02341206), ACTB (Hs03023943), GAPDH (Hs02758991), IGFBP1 (Hs00236877), PRL (Hs00168730). A plate standardization control comprising pooled primary placenta and decidua RNA was analyzed for its expression for all genes examined to allow combination of results from separate plates. All samples (including plate standardization control) were run in triplicate. Gene expression was quantified using Bio-Rad CFX Manager Software version 3.1. Samples from individual donors were run separately, normalized by both GAPDH and ACTB expression, further normalized by the plate standardization control, and their results were then combined for display on the plots shown.

### Flow cytometry

Flow cytometry was performed on freshly thawed and washed primary dNK cells. Prior to staining, a portion of dNK cells from each sample was pooled together and used to create Live/Dead, unstained, and fluorescence-minus-one controls. Live/Dead staining was performed using Live/Dead Fixable Violet stain (Invitrogen L34964), with the Live/Dead control prepared by snap freezing half of its cells to ensure sufficient cell death. Details of all antibodies used and their dilutions are described in Supplementary Table 2. Each sample and control was prepared in PBS containing 1% BSA. Samples were first treated with Human TruStain FcX (BioLegend 422302) diluted 1:20 in PBS containing 1% BSA for 20 minutes. Samples were then washed and stained with fluorophore-conjugated antibodies for one hour at room temperature in opaque tubes. Samples were then washed, fixed in formalin for 10 minutes, and washed again. Compensation controls were prepared for each antibody using UltraComp eBeads Compensation Beads (Invitrogen 01-2222). Samples were then analysed with the BD LSRFortessa X-20 flow cytometer at the Temerty Faculty of Medicine Flow Cytometry Facility, University of Toronto.

### Immunohistochemistry and cavity measurement

Cultures were removed from Cultrex gel domes using 500 μL Organoid Recovery Solution (Cultrex, R&D #3700) per dome and incubating plates for 75 minutes at 4°C with intermittent agitation. Recovered samples were washed and fixed for 15 minutes in formalin. After fixation, samples were washed and placed into 75% ethanol prior to embedding. Embedding was performed at the University of Toronto’s Microscopy Imaging Laboratory. Samples were recovered from ethanol and then spun in low melting point agar to aggregate organoids, all of which were then embedded in paraffin blocks. Blocks were cut into 10μm sections using a microtome. Serial sections were performed and multiple sections were stained in parallel on each slide to ensure that all staining and analysis was performed near the middle of the organoids. Slides were dewaxed using two 10-minute incubations in xylene, and dehydrated through two 5-minute incubations each of 100% and 95% ethanol. Slides were then incubated in peroxide for 10 minutes, followed by antigen retrieval using pH 6 sodium citrate with a vegetable steamer for all antibodies. Details of all antibodies used and their dilutions are described in Supplementary Table 2.

All primary antibodies were prepared in Blocker Casein (Thermo Scientific 37532) with 1% Tween 20 (BioShop TWN510) which were applied after slides had cooled following antigen retrieval. Slides were incubated with primary antibodies overnight at 4°C in a humidified chamber. Controls slides which were incubated in diluent without any primary antibodies were prepared during each experiment to ensure specificity of staining. Slides were then washed and incubated for 30 minutes with secondary antibodies at room temperature. Antibody staining was visualized using SignalStain DAB Substrate Kit (Cell Signaling Technology #8059S). Slides were counterstained with hematoxylin (Cell Signaling Technology 14166S) for two minutes, and dipped 7 times in 0.1% sodium bicarbonate for bluing. Slides were then cleared and dehydrated by dipping 15 times each in two buckets of 95% ethanol, two buckets of 100% ethanol, and two buckets of xylenes. Slides were mounted and coverslipped using Permount media (Fisher Chemical SP15). Slide scans were obtained using the Zeiss Axioscan slide scanner at the University of Toronto’s Microscopy Imaging Laboratory. DAB staining quantification was performed using QuPath Software version 0.4.4 thresholder and cell detection tools by two different operators who were blinded to whether each sample contained mono- or co-cultured organoids. Measurement of the proportion of organoid area occupied by cavities was performed using QuPath Software version 0.4.4 pixel classification tools. Cavity measurement was performed on the same images obtained from organoids stained for SDC1, a marker which borders the STB which typically surrounds cavities.

### RNA isolation and sequencing

All samples were collected and snap frozen following seven days of culture, then processed according to the manufacturer’s instructions using the RNeasy Micro Kit (Qiagen 74004) for total RNA isolation. The sample disruption and homogenization steps were performed using QIAShredders (Qiagen 79654). Libraries were prepared at the Donnelley Centre at the University of Toronto using Takara SMART-Seq v.4 Ultra Low Input RNA kits. Sequencing was also performed at the same facility using the NovaSeq 6000 SP to generate paired-end reads with a target of 35 million reads per sample. There was heterogeneity in methods of sample preparation due to attempts to separate culture constituents after co-culture using a Ficoll gradient for some, but not all samples. After sequencing, the expression was not found to differ significantly between samples that were immediately snap frozen after culture and those that were placed on a Ficoll gradient before snap freezing when comparing replicate cultures that contained the same constituents. Reads from samples prepared through these heterogeneous methods were then combined using batch correction.

FASTQ files were quality assessed by FASTQC, filtered for low quality reads and adapter sequences using cut adapt. Cleaned reads were aligned using HISAT2 (2.2.1), against the human genome build GRCH37. Sam files were converted to BAM files, sorted and indexed using samtools (1.13, htlib 1.13+ds). R (4.1) scripts were used to create a count table from BAM files and the human transcriptome G (including coding and non-coding RNA species) using the featureCounts function of Rsubread (2.8.2). Differential expression was calculated using a pipeline of EdgeR (4.2.1), and Limma (3.60.3) packages. Gene set enrichment was calculated by the camera function in the limma package. Human gene ontology files were obtained from the enrichment map resource (download.baderlab.org/EM_Genesets/). Human placental ontologies were from supplemental files from Naismith and Cox, 2021. Barcode graphs were generated using the barcode function in limma.

Removal of dNK gene expression was accomplished using the granulator package in R applying the detanglr algorithm to estimate the transcriptional contribution of the NK cells to the co-cultured bulk mRNA signal. The dNK and organoid samples were used to estimate the pure signal. The estimated mean contribution was removed in limma as a fitted model.

Data sets are deposited at GEO under the accession number GSE272695.

### Masson’s Trichrome

Slides were dewaxed using two 10-minute incubations in xylene, two 5-minute incubations in 100% ethanol, and two 5-minute incubations in 95% ethanol, followed by rehydration through one 5-minute incubation in distilled water. Slides were then re-fixed in Bouin’s solution (MilliporeSigma HT10132) overnight at room temperature, rinsed in distilled water, and washed thoroughly in tap water. Next, slides were stained with Weigert’s hematoxylin (MilliporeSigma 1159730002) for 5 minutes, washed in tap water for 5 minutes, and stained with Biebrich Scarlet-Acid Fuchsin for 10 minutes (MilliporeSigma HT151). Slides were then briefly rinsed in distilled water and differentiated in 5% phosphomolybdic-phosphotungstic acid solution for 10 minutes (MilliporeSigma 221856 and 79690 in distilled water), directly followed by incubation in Aniline blue (MilliporeSigma B8563) for 5 minutes. Slides were then rinsed in distilled water, developed in 1% glacial acetic acid solution for 5 minutes, cleared through single dips in two buckets each of 95% and 100% ethanol, and prepared for mounting by dipping 15 times in two buckets of xylenes. Slides were mounted and coverslipped using Permount media (Fisher Chemical SP15). Images were analysed using QuPath Software version 0.4.4 pixel classification and point counting tools.

### Secreted factor measurement

Culture media from days two and seven of mono- and co-cultures was submitted to Eve Technologies for analysis through multiplex laser bead-based panels. Two sets of samples were submitted for different analyses as described in Supplementary Table 1. The first set of samples was subjected to the Human Cytokine/Chemokine 48-plex Discovery Assay Array (Eve Technologies HD48). The second set of samples was subjected to both the Human MMP and TIMP Discovery Assay Array (Eve Technologies HMMP/TIMP-C,O) and the Human Angiogenesis and Growth Factor 17-Plex Discovery Assay Array (Eve Technologies HDAGP17) in parallel.

### Statistical analysis

All statistical analysis was performed by custom R scripts running in R Studio version 2022.12.0+353. For comparisons between mono- versus co-cultured organoids that were made for one continuous variable at a time, either unpaired two-tailed t-tests or Wilcoxon rank sum tests with continuity correction were performed as indicated in Supplementary Table 1. Shapiro-Wilk normality tests were performed to determine which statistical test to use: data which did not follow a normal distribution with Shapiro p<0.05 was assessed with a Wilcoxon rank sum test, and data which did follow a normal distribution with Shapiro p>0.05 was assessed with a t-test. For comparison between mono- versus co-cultured organoids for categorical variables, Pearson’s Chi-squared test with Yates’ continuity correction was performed. For multi-variable comparisons between mono- and co-cultured organoids and dNKs (used for analysing secreted factor panels), one-way ANOVAs with Tukey’s Honestly Significant Differences tests were performed.

Transcriptomic analysis used linear models in limma with a false discovery correct p-value accepting <0.05 as significant. For gene set enrichment a false discovery corrected p-value of <0.05 was also consider significant, however it should be noted that this is often viewed are overly stringent.

## ACKNOWLEDGEMENTS

We thank our core facilities managers and staff at The University of Toronto Temerty Faculty of Medicine including Lindsey Fiddes and Nathaniel Windsor at the Microscopy Imaging Laboratory, Nathalie Simard at the Flow Cytometry Facility, and Sherin Mohammed Shibin at the Donnelley Centre. Thank you to Pascale Robineau-Charette and Andrea Jurisicova for your time discussing this work.

## DATA AVAILABILITY

GSE272695

