## Supplemental Figures and Tables for "Decidual natural killer cells promote extravillous trophoblast developmental pathways: evidence from trophoblast organoid co-cultures"

Table S1: Sample sizes, composition, and statistical tests:

| Experiment | Org donors | dNK donors | Org+dNK combinations | Mono-cultured org | Co-cultured org | Statistical test |
|---|---|---|---|---|---|---|
| 2D area comparison | 4 | 5 | 9 | 50 | 82 | t-test |
| Organoid counts | 6 | 7 | 13 | 2943 | 2731 | t-test |
| Cavity comparison | 4 | 4 | 6 | 34 | 54 | Wilcoxon test |
| Ki67 staining | 4 | 5 | 7 | 55 | 73 | Wilcoxon test |
| Bulk RNA sequencing | 2 | 2 | 4 | N/A | N/A | Linear models with FDR-corrected p-value |
| CD44 staining | 5 | 5 | 8 | 49 | 67 | Chi-squared test |
| Masson's trichrome - area | 5 | 7 | 10 | 64 | 55 | Wilcoxon test |
| Masson's trichrome – point counts | 5 | 7 | 11 | 108 | 84 | Wilcoxon test |
| E-cadherin staining | 4 | 3 | 6 | 21 | 20 | t-test |
| SDC1 staining | 4 | 4 | 6 | 34 | 54 | t-test |
| Cytokine and chemokine measurement | 2 | 3 | 6 | N/A | N/A | ANOVA |
| Growth/ECM-modifying factor measurement | 3 | 2 | 6 | N/A | N/A | ANOVA |

Table S2: Antibody details:

| Target | Species of origin | Catalogue number | Company | Application | Dilution |
|---|---|---|---|---|---|
| Cytokeratin 7 | Mouse | M7018 | Dako | ICC | 1 in 200 |
| GATA3 | Goat | AF2605 | R&D | ICC | 1 in 50 |

| | | | | | |
|---|---|---|---|---|---|
| Anti-mouse-AF488 | Donkey | A21202 | Life Technologies | ICC | 1 in 500 |
| Anti-goat-AF647 | Donkey | A21447 | Invitrogen | ICC | 1 in 500 |
| E-cadherin | Rabbit | 3195 | Cell Signalling | IHC | 1 in 2000, DAB for 3 minutes |
| Syndecan-1 | Goat | AF2780 | R&D | IHC | 1 in 100, DAB for 2.5 minutes |
| Ki67 | Rabbit | Ab16667 | Abcam | IHC | 1 in 400, DAB for 7 minutes |
| Anti-Rabbit SignalStain Boost HRP | Goat | 8114S | Cell Signalling | IHC | Not diluted |
| Anti-Goat SignalStain Boost HRP | Horse | 63707S | Cell Signalling | IHC | Not diluted |
| CD56-PerCP Cy5.5 | Mouse | 392420 | Biolegend | FC | 1 in 20 |
| CD45-AF647 | Mouse | 393406 | Biolegend | FC | 1 in 20 |

ICC= immunocytochemistry (fluorescent), IHC= immunohistochemistry (chromogenic), FC= flow cytometry

A

| Gestational age (weeks) | % CD56+ | % CD45+CD56+ |
|---|---|---|
| 5.5 | 92.0 | 76.7 |
| 6.2 | 87.6 | 44.3 |
| 6.3 | 84.1 | 53.4 |
| 6.5 | 95.3 | 88.5 |
| 7.5 | 92.3 | 73.6 |
| 8.2 | 95.2 | 41.9 |
| 9.0 | 97.0 | 90.7 |
| 11.0 | 94.6 | 80.9 |
| 11.0 | 93.0 | 78.9 |
| 11.1 | 97.0 | 88.2 |
| Concatenated | 91.1 | 65.4 |

B

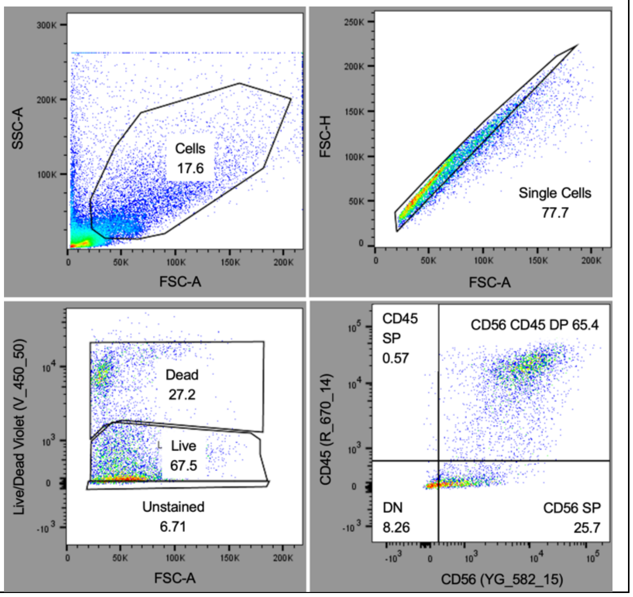

Fig. S1. **dNK analysis details. A.** Proportions of CD56+ and CD45+ cells after dNK enrichment from ten individual donors assessed by flow cytometry. **B.** Gating strategy used for analysis of all flow cytometry samples, shown with concatenated samples. Populations were gated by selecting for cells (excluding debris), single cells, live cells, and then analyzing CD56 and CD45 staining, in that order.

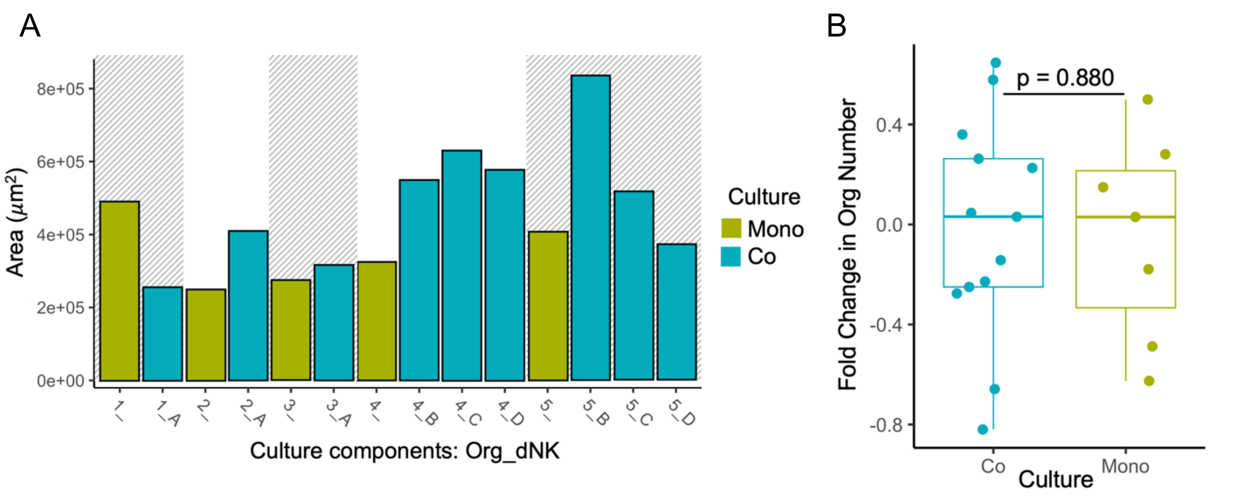


Fig. S2. **Organoid area of individual co-cultures and organoid counts. A.** Most combinations of organoid and dNK donors show increased 2D area of organoids after 7 days. Trophoblast organoid (Org) donors are denoted with numbers 1-5, and dNK donors are denoted with letters A-D, with conditions described following the format of "Org_dNK". Trophoblast organoids from the same donor are visually grouped by the presence or absence of background shading. **B.** No change in organoid numbers occurs after co-culture with dNKs for 7 days (p=0.90). As standardization of organoid numbers at the outset of culture was not possible, organoid number is represented as fold change between day 0 and day 7 of culture for comparison.

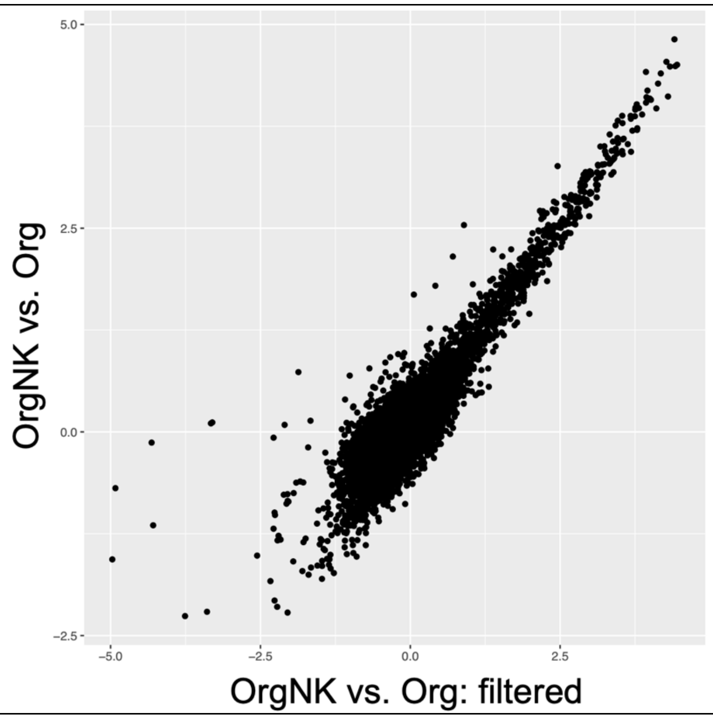


Fig. S3. **dNK signal removal from co-cultured organoid gene expression.** Filtering removes dNK-associated expression (observed in dNK monocultures) is removed from analysis of co-cultured organoids. After this step, dNK signals do not contribute to measured differential expression between trophoblast organoid co-cultures versus monocultures.

A

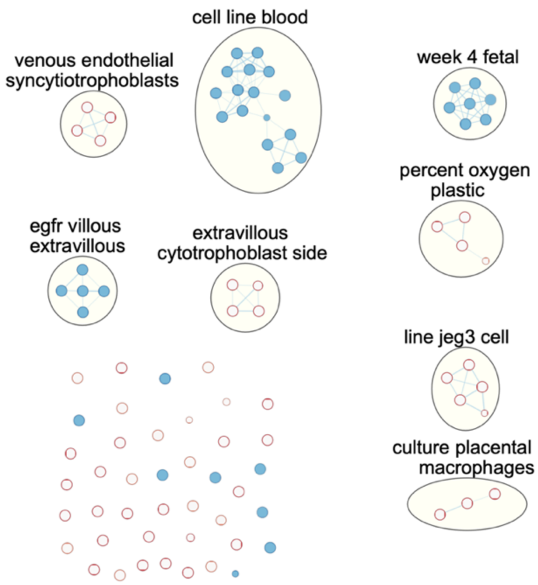


B

| | HLA-G | CD44 | ASCL2 | FOXO1 | ITGA1 | ITGA2 |
|---|---|---|---|---|---|---|
| OrgNK vs. Org Expression | 0.55 | 3.33 | 1.14 | 3.1 | 2.4 | -0.34 |
| Org vs. NK Expression | 3.39 | -6.67 | 5.87 | -5.75 | -5.85 | 0.06 |
| OrgNK vs. NK Expression | 3.67 | -3.02 | 6.35 | -2.72 | -3.02 | -0.01 |
| p-value | 0.21 | <0.001 | <0.001 | 0.02 | <0.001 | 0.18 |

Fig. S4. **Trophoblast subtype gene expression. A.** Placental gene sets of co-cultured organoids visualized with Cytoscape. Placental gene sets observed in co-cultured organoids mostly relate to EVT differentiation. **B.** Expression changes in select EVT-associated genes. While HLA-G, the canonical EVT marker, and ITGA2, a column CTB marker, did not show statistically significant changes in expression, other EVT markers (CD44, ASCL2, FOXO1, and ITGA1) did show significant upregulation in co-cultured organoids.

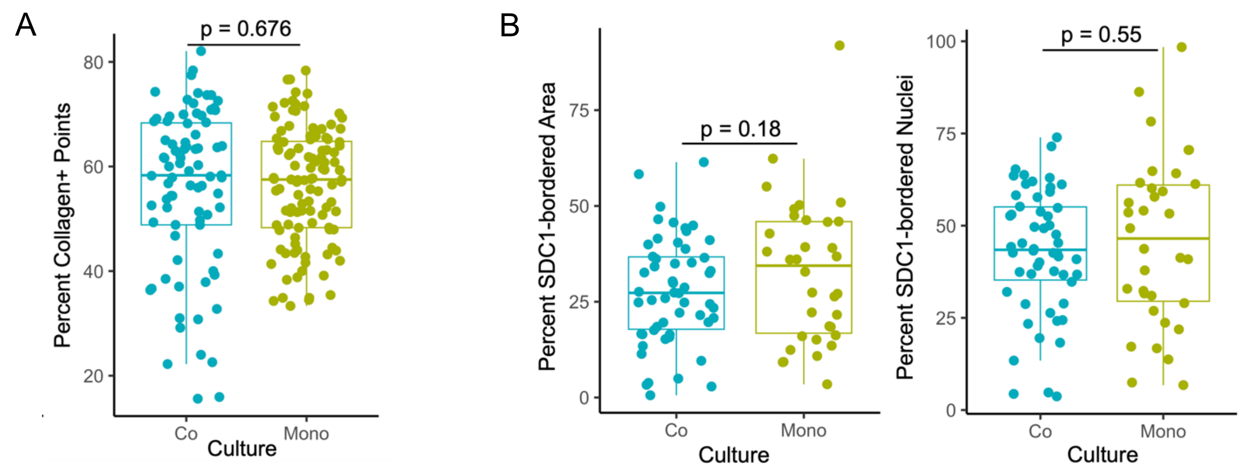


Fig. S5. **Additional histological measurements. A.** Point counting of Masson's trichrome-stained mono- and co-cultured organoids reveals that there was no difference in amount of collagen-positive points within organoids, and thus no change in distribution of collagen within organoids. **B.** Syncytiotrophoblast in organoid mono- and co-cultures. No change in proportion of organoid area bordered by SDC1+ membrane or in number of nuclei bordered by SDC1+ membrane in mono- versus co-cultured trophoblast organoids.

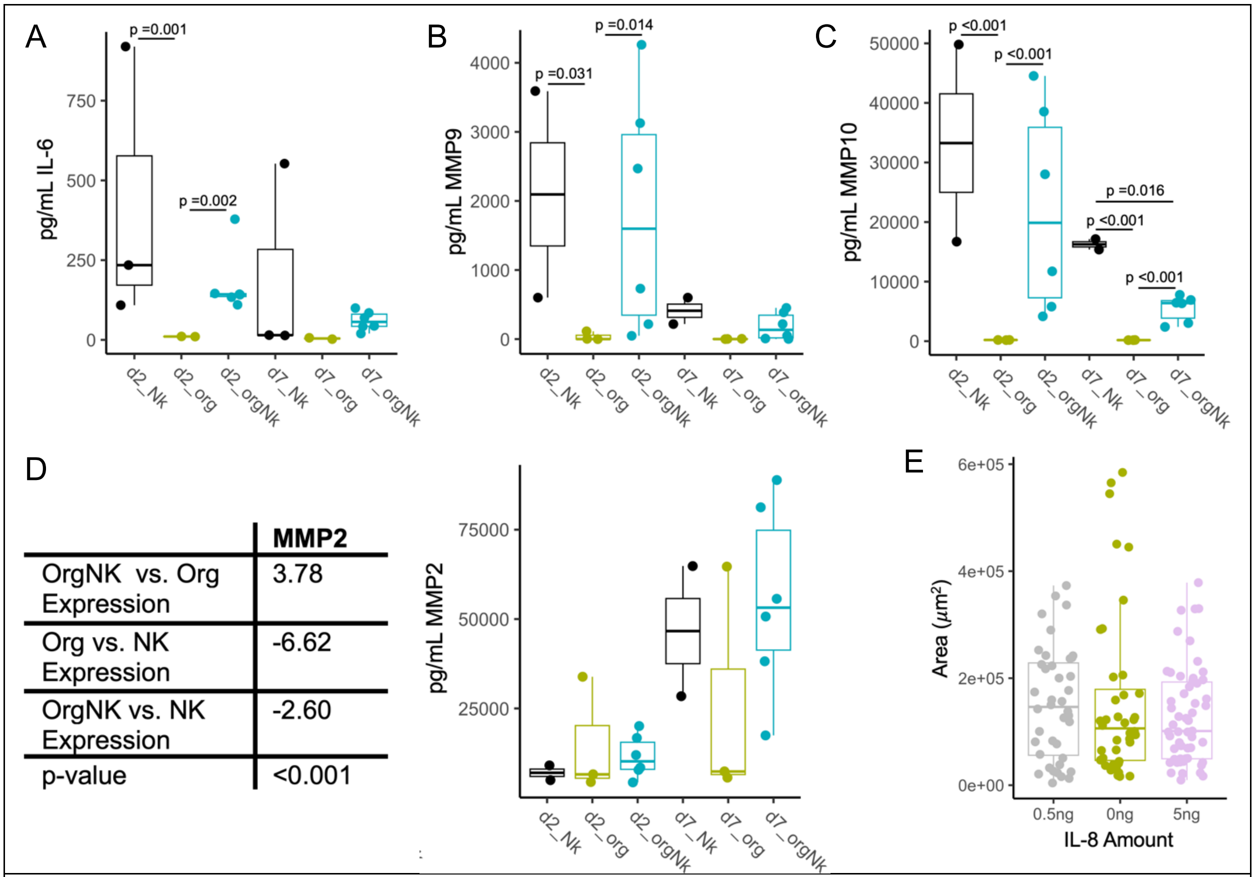


| | MMP2 |
|---|---|
| OrgNK vs. Org Expression | 3.78 |
| Org vs. NK Expression | -6.62 |
| OrgNK vs. NK Expression | -2.60 |
| p-value | <0.001 |

Fig. S6. **Additional secreted factors A.** IL-6 **B.** MMP9 **C.** MMP10. **D.** Significant differential expression of the MMP2 gene was observed between co-cultured and monocultured trophoblast organoids in transcriptomic analysis. No significant differences were observed between amounts of MMP2 found in culture media of monocultured trophoblast organoids, monocultured dNKs, or their co-cultures. **E.** IL-8 addition to culture media did not impact trophoblast organoid size in the absence of dNKs.

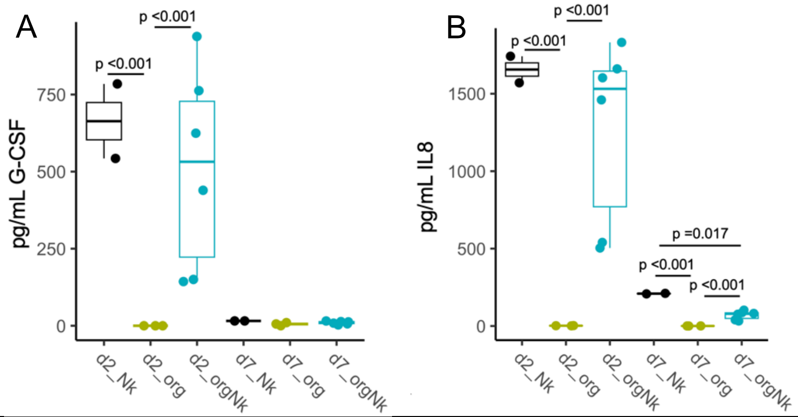


Fig. S7. **Repeat analysis of IL-8 and G-CSF levels in culture supernatants. A.** G-CSF and **B.** IL-8 levels were measured alongside extracellular matrix-modifying and angiogenic factors in the 30-plex panel (data presented for G-CSF and IL-8 in the main text was obtained from the 48-plex panel of immunomodulatory factors). The same trends in levels of these factors were observed in this separate analysis performed on different samples.
